# “Like Sugar in Milk”: Reconstructing the genetic history of the Parsi population

**DOI:** 10.1101/128777

**Authors:** Gyaneshwer Chaubey, Qasim Ayub, Niraj Rai, Satya Prakash, Veena Mushrif-Tripathy, Massimo Mezzavilla, Ajai Kumar Pathak, Rakesh Tamang, Sadaf Firasat, Maere Reidla, Monika Karmin, Deepa Selvi-Rani, Alla G. Reddy, Jüri Parik, Ene Metspalu, Siiri Rootsi, Kurush Dalal, Shagufta Khaliq, Syed Qasim Mehdi, Lalji Singh, Mait Metspalu, Toomas Kivisild, Chris Tyler-Smith, Richard Villems, Kumarasamy Thangaraj

## Abstract

**Background:** The Parsis, one of the smallest religious community in the world, reside in South Asia. Previous genetic studies on them, although based on low resolution markers, reported both Iranian and Indian ancestries. To understand the population structure and demographic history of this group in more detail, we analyzed Indian and Pakistani Parsi populations using high-resolution autosomal and uniparental (Y-chromosomal and mitochondrial DNA) markers. Additionally, we also assayed 108 mitochondrial DNA markers among 21 ancient Parsi DNA samples excavated from Sanjan, in present day Gujarat, the place of their original settlement in India.

**Results:** Our extensive analyses indicated that among present-day populations, the Parsis are genetically closest to Middle Eastern (Iranian and the Caucasus) populations rather than their South Asian neighbors. They also share the highest number of haplotypes with present-day Iranians and we estimate that the admixture of the Parsis with Indian populations occurred ∼1,200 years ago. Enriched homozygosity in the Parsi reflects their recent isolation and inbreeding. We also observed 48% South-Asian-specific mitochondrial lineages among the ancient samples, which might have resulted from the assimilation of local females during the initial settlement.

**Conclusions:** We show that the Parsis are genetically closest to the Neolithic Iranians, followed by present-day Middle Eastern populations rather than those in South Asia and provide evidence of sex-specific admixture from South Asians to the Parsis. Our results are consistent with the historically-recorded migration of the Parsi populations to South Asia in the 7thcentury and in agreement with their assimilation into the Indian sub-continent’s population and cultural milieu “like sugar in milk”. Moreover, in a wider context our results suggest a major demographic transition in West Asia due to Islamic-conquest.

## Background

The Parsi (or Parsee) community of the Indian sub-continent are a group of Indo-European speakers and adherents of the Zoroastrian faith, one of the earliest monotheisms that flourished in pre-Islamic Persia (present-day Iran)(1,2). Zoroastrianism was the official religion of Persia from 600 B.C. to 650 A.D.(3-5) and despite a long history of well-preserved culture, it now has a limited number of followers (6,7). The Heritage Institute estimates the total number of Zoroastrians to be around 137,000, with 69,000 living in India, roughly 20,000 in Iran (8) and 2,000-5,000 in Pakistan (8,9). This reduction in population is mainly due to strict marriage practices and low birth rates (7,10-12).

The Parsi trace their ancestry to the Zoroastrians of modern Iran who are followers of the Prophet Zoroster or Zarathushtra (3). In the seventh century the Zoroastrian Sassanian dynasty was threatened by Islamic conquest and a small group of Zoroastrians fled to Gujarat in present-day India, where they were called ‘Parsi’ (literally meaning ‘people from Paras or Faras’, the local term for Persia)(3,4,6,13). Several myths narrate their first arrival in the West Coast of India and settlement in Sanjan (Gujarat) (4,14-17). The most popular one mentioned in the *Qissa-e-Sanjaan* is that an Indian ruler called *Jadi Rana* sent a glass full of milk to the Parsi group seeking asylum (4,18). His message was that his kingdom was full with local people. The Zoroastrian immigrants put sugar (or a ring, in some versions of the story) into the milk to indicate an assimilation of their people into the local society, like “sugar in milk”(14,18) In contemporary India and Pakistan, we see their adoption of local languages (Gujarati, Sindhi) and economic integration while maintaining their ethnic identity and practicing strict endogamy (1,3,4,12,19-21).

Previous genetic analyses of the Parsis have focussed mainly on low-resolution uni-parental markers which have suggested their affinity with both West Eurasian and Indian populations(22-24). Autosomal analysis based on microsatellites or human leukocyte antigens (HLA) have revealed their intermediate position among the populations of South Asia and the Middle East/Europe(9,24). A study of mitochondrial DNA (mtDNA) variation reported 60% of South Asia lineages among the Pakistani Parsi population (23), whereas the male lineages based on Y chromosome admixture estimates were almost exclusively Iranian (22). Based on these results, a male-mediated migration followed by assimilation of local South Asia females was concluded (23).

These early studies of the Parsi populations relied mainly on low-resolution markers, limiting the power of the analyses (9,23-25), and the majority of Parsis (∼ 98.8%), living in India, have been underrepresented in these studies. In the present study, we used a combination of high-resolution uni-parental and bi-parental genetic markers on modern samples, as well as mtDNA markers on ancient Parsi samples excavated from Sanjan, in present-day Gujarat, India (Fig. 1). The human remains from Sanjan *dokhama* (tower of silence) District Valsad, Gujarat, was excavated in 2004. The AMS (Accelerator Mass Spectrometry) dating of human remains suggest that the *dokhama* belongs to the 14th to 15th century A.D. (17).

**Fig. 1.**
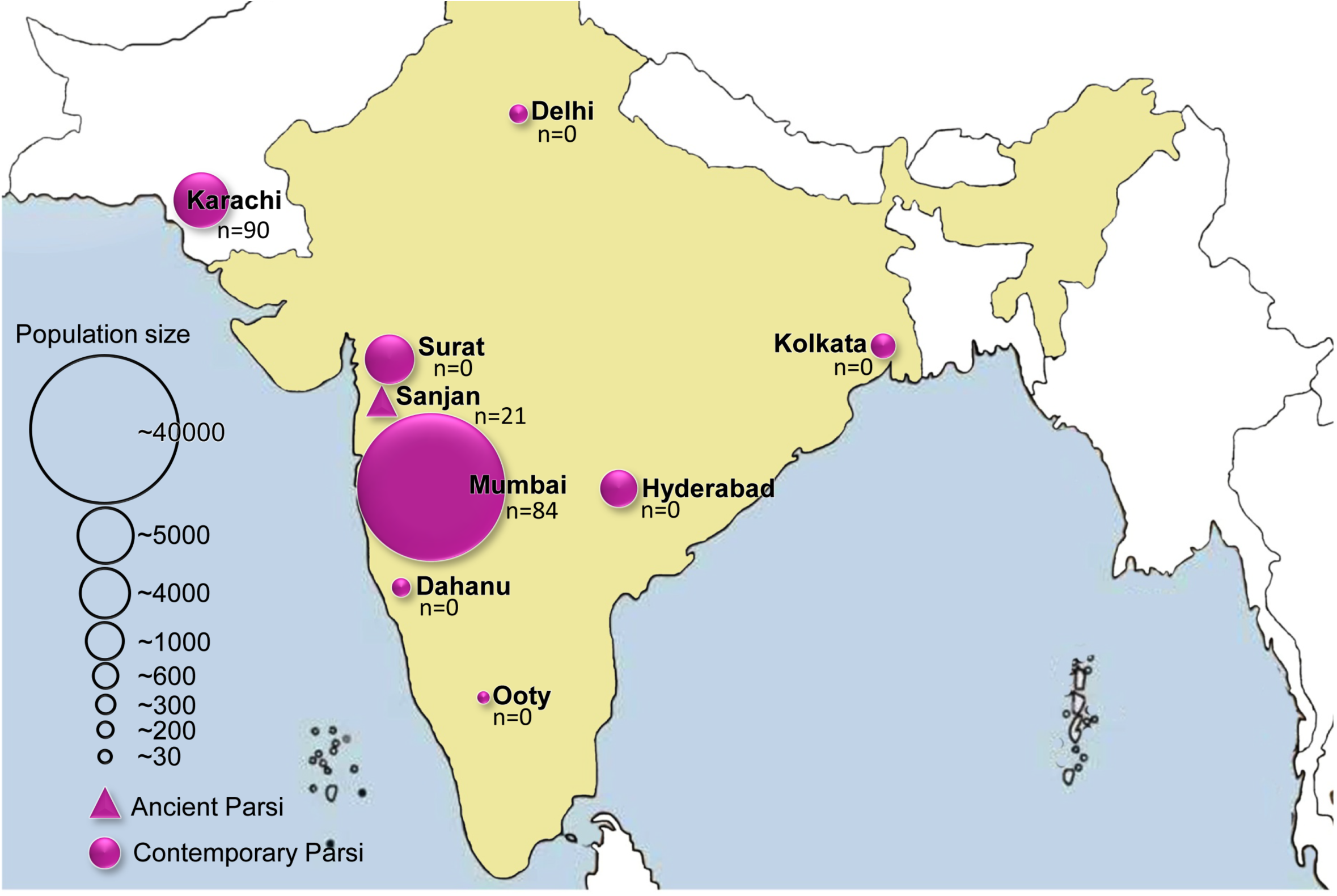
The geographical distribution and sampling locations of modern and ancient Parsi samples. The population data is obtained from Parzor foundation, New Delhi, India.

We investigated whether or not the current Parsi people living in India and Pakistan are genetically related amongst themselves and with the present-day Iranian population, and if their genetic composition has been affected by the neighbouring Indian and Pakistani populations. We also examined runs of homozygosity (RoH) to study consanguinity. To address the extent to which the current Parsi populations assimilated local females during their long formation history, we compared their mtDNA haplogroup composition with ancient remains excavated in Sanjan, the initial settlement established by these migrants from Persia (17).

## Results and Discussion

For autosomal analyses, we used Illumina HumanHap 650K genotyping chips on 19 Indian Parsi samples collected from Mumbai, and Illumina 2.5M genotyping chips for 24 Pakistani Parsi individuals from Karachi (Fig. 1 and Additional File 1). The combined Parsi dataset was merged with a global data from the published literature (26,27). Table S2 lists the populations and number of SNPs used for various analyses after quality control (Additional File 1). The mean allele frequency differentiation between the two Parsi (Indian and Pakistani) groups was the lowest (*F*_ST_ Indian and Pakistani Parsi = 0.00033 ± 0.00025), followed by the differentiation of each from the Iranian population (0.011 ± 0.00021 and 0.012 ± 0.00025 for Pakistani and Indian Parsis, respectively), suggesting a common stock for both the Indian and Pakistani Parsis with the closest interpopulation affinity with populations from their putative homeland, Iran (Additional File 1: Fig. S1 and Additional File 1: Table S3). Collectively, in *F*_ST_-based analysis, both of the Parsi groups showed significantly closer connection with West Eurasians than any of the Indian groups (two-tailed p value <0.0001).

We applied the default settings of the SmartPca programme implemented in the EIGENSOFT package (28) and performed principal-components analysis (PCA) with other Eurasian populations using autosomal SNP data (Additional File 1: Table S2 and Supplementary text). Our plot of the first and second principal components (PCs) clusters the Indian and Pakistani Parsis together, along the European-South Asian cline (Fig. 2). A plot of populationwise mean of eigenvalues showed their placement between the Pakistani and Iranian populations, indicating that the Parsis might have admixed from these two groups. Such an intermediate position of Parsis closer to Iranians than to their present geographic neighbours (Sindhi and Gujarati), suggests that the Parsis may have major ancestry from West Eurasians (Iranians) and minor ancestry with South Asians (Fig. 2 and Additional File: Fig. S1). In order to identify the ancestry components of Parsis more quantitatively, we applied the model-based clustering method ADMIXTURE (29). The best model (30,31) suggested eight major ancestral groups and identified three ancestral components within the Parsi (Fig. 3 and Additional File 1: Table S4). The distributions of these components among Indian and Pakistani Parsis were unique and similar to each other, but were distinct from their neighbouring South Asian populations (Additional File 1: Fig. S2a). The ANI (Ancestral North Indian) ancestry calculated from *f*4 ancestry estimate showed a substantially higher level of this ancestry among Parsis than any other South Asia population (Additional File 1: Fig. S2b). Supporting the *F*_ST_ and PCA results, the Middle-Eastern-specific (blue) ancestry component was significantly higher (two-tailed p value <0.0001), in the Parsis than in any other populations residing in South Asia that were examined. The present-day Iranian population exhibited a striking difference from the Parsis, mainly in carrying an additional European component (light blue) and substantially lower South Asian ancestry (dark green) (Fig. 3 and Additional File 1: Fig. S2 and Table S4). It was suggested previously that the Islamic conquest had a major genomic impact on several Middle Eastern populations, including Iranians (32). Since Parsis diverged from Iranians just after this conquest, they may represent the genetic strata of Iran before Islamic-conquest. To test this scenario, we applied a formal test of admixture *f*3 statistics (Additional File 1: Table S5). For Iranians, a negative value with significant Z-scores indicated that they are descendants of a population formed by the admixture of Neolithic Iranians and populations from the Arabian-Peninsula, while for Parsis this test was positive with significant Z-scores. Therefore, it seems plausible that the additional light blue component may have been introduced to Iran after the exile of the Parsis, likely via recent gene flow from the Arabian Peninsula(32-34). The quantitative estimation of South Indian and Iranian ancestries among Parsis and their South Asian neighbours showed a significant level of differentiation in ancestry composition with an inclination of Parsis towards Iranian ancestry (two-tailed p value <0.0001) (Table 1 and Additional File 1: Supplementary text).

**Fig. 2.**
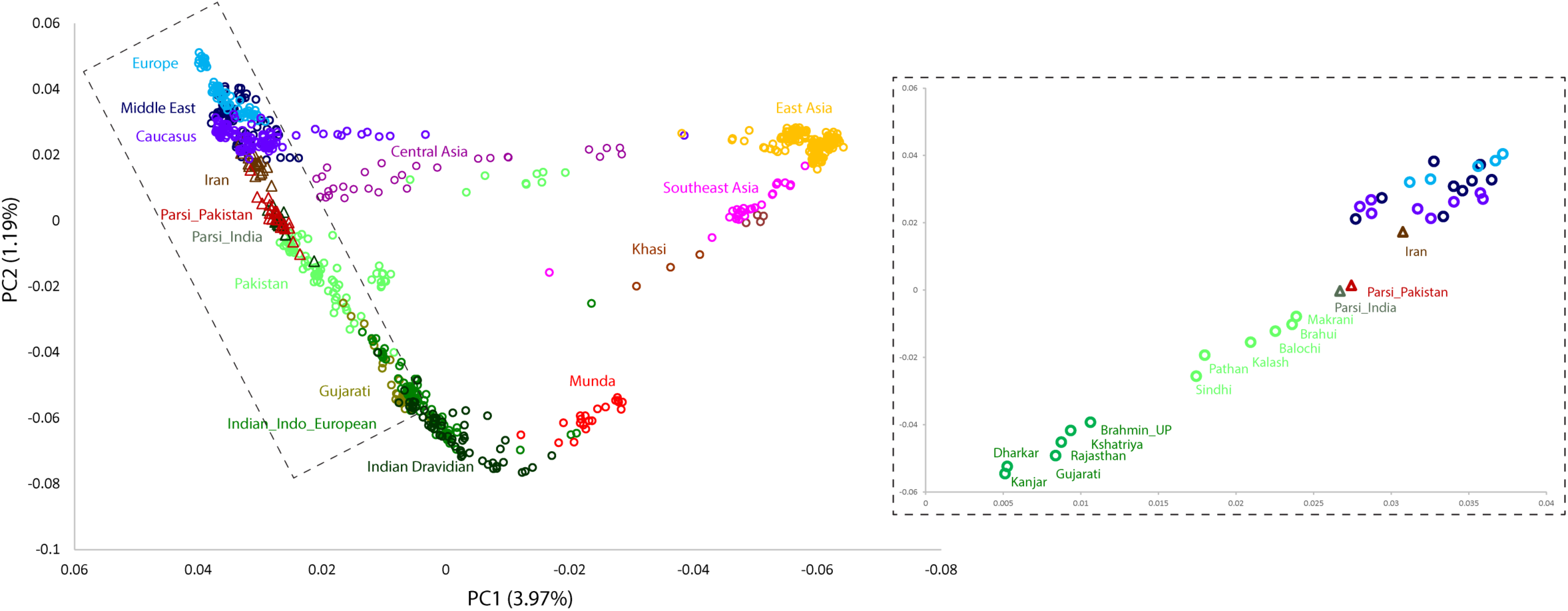
Principal component analysis of the combined autosomal genotypic data of individuals from Eurasia. The inset shows a plot of mean eigenvalues of Parsi and other closeby populations.

**Table 1.** The South Indian (SIND), and Iranian ancestry among Parsis and neighbouring populations

| Population (X) | SIND (SE) | Iranian (SE) |
| --- | --- | --- |
| Pathan | 31 (2.4) | 56.5 (1.8) |
| Sindhi | 22.9 (2.7) | 63.1 (1.9) |
| Parsi_Pakistan | 6.4 (2.4) | 76.6 (1.5) |
| Parsi_India | 8.8 (2.5) | 74.6 (1.7) |
| Gujarati | 58.1 (2) | 34.3 (1.6) |

We computed *D* statistics (35) to determine the nature of gene flow and admixture of Parsis with their parental (Iranian) and neighbouring (Gujarati, Sindhi) populations (Table 2). Consistent with the previous analyses (Figs. 2 and 3), both of the Parsi populations shared a highly significant value of *D* with each other. On the other hand, the South Asian populations (Gujarati and Sindhi) had significant levels of gene flow with each other, as well as with both of the Parsi populations when evaluated with respect to the present-day Iranian population (Table 2). Two independent studies have recently reported the data from ancient Iranian samples (36,37). It was suggested that the early Neolithic Zagros sample showed closer affinity with the Iranian Zoroastrians (36). Here we estimated the *D* values of Parsis for Neolithic Iranians *vs* modern Iranians to compare the allele sharing. Our results demonstrated a significant level of genetic affinity between Parsis and Neolithic Iranians (Table 2 and Additional File 1: Supplementary text and Table S6). The outgroup *f*3 statistics of ancient Iranian samples supported the close affinity of the Parsis with the Neolithic Iranians (Additional File 1: Supplementary text, Fig. S3 and Table S6). Moreover, for modern populations, the outgroup *f*3 statistic test and identity-by-state (IBS) plots supported the closer affinity of the Parsis with the West Eurasian populations than South Asians (Additional File 1: Figs. S4 and S5). In order to compare the shared drift with Iranian and Indian (South Munda) populations, we plotted the derived allele sharing values of Parsis and other Eurasian populations calculated with respect to the Iranian and South Munda (Indian) populations (Additional File 1: Supplementary text and Fig. S6). This analysis aligned Parsis closer to the Iranian axis between Pakistani and West Eurasian populations, confirming the historical interpretation of the most recent common ancestry of Parsis with the Iranians. TreeMix (38) analysis support these conclusions and shows the Parsis located between the South Asian and Iranians (Additional File 1: Fig. S7).

**Fig. 3.**
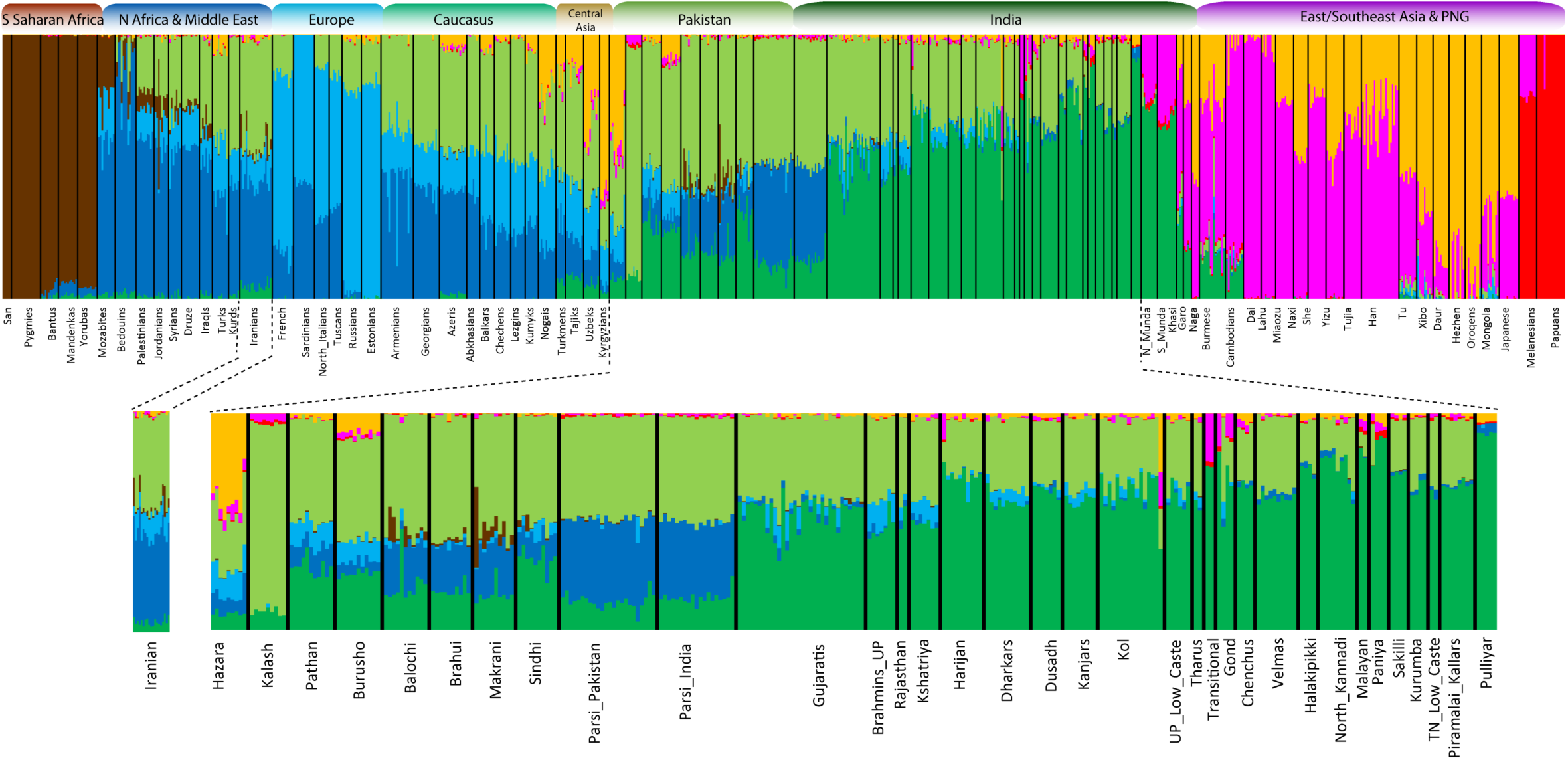
Results of ADMIXTURE analysis (*k* = 8) of world populations with a zoom-in on Iranian, Parsis and other South Asian populations.

**Table 2.**
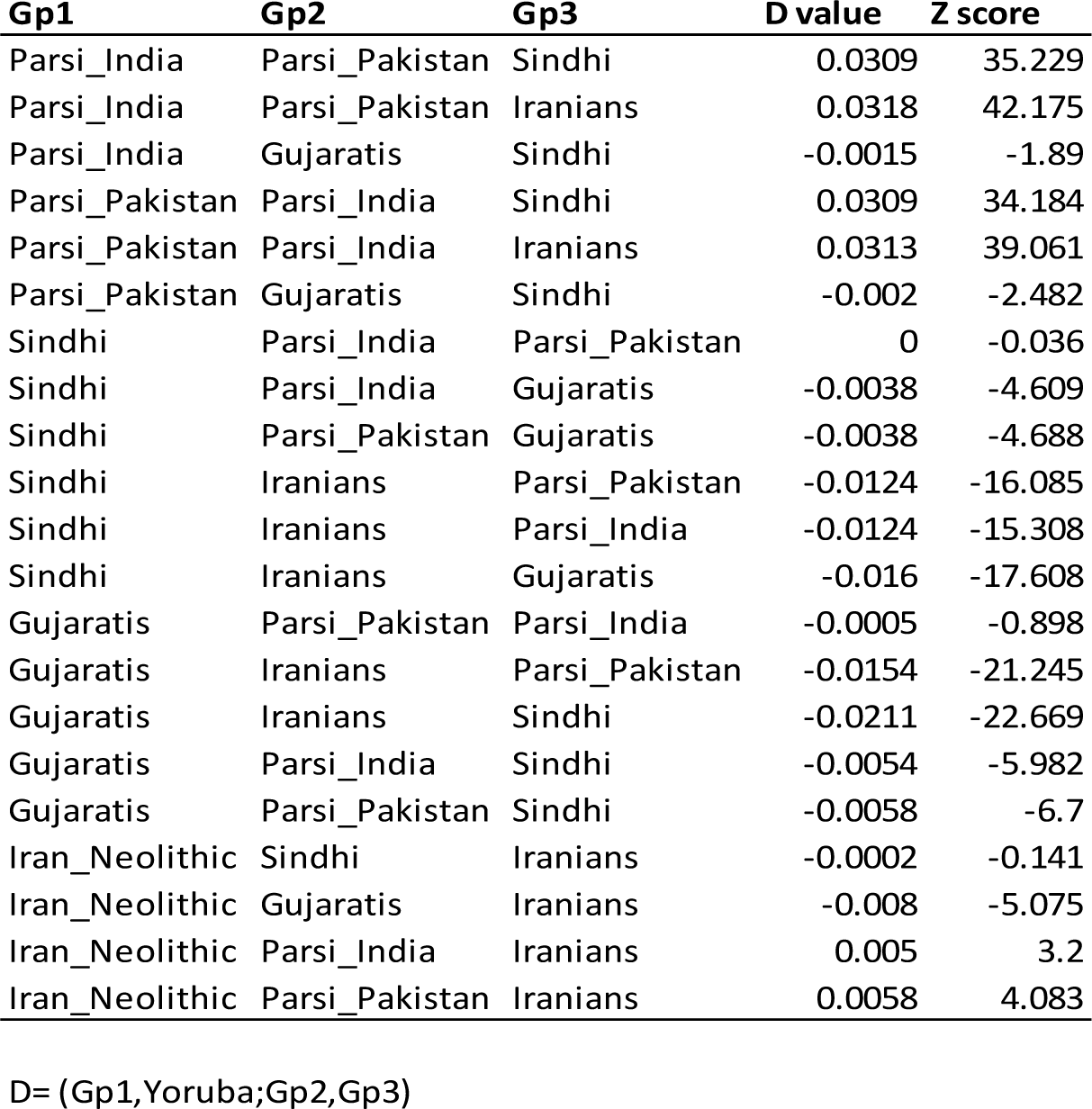
The test of geneflow (*D* statistics) between Parsis, modern Iranian, Neolithic Iranian, Sindhi and Gujarati

| Gp1 | Gp2 | Gp3 | D value | Z score |
| --- | --- | --- | --- | --- |
| Parsi_India | Parsi_Pakistan | Sindhi | 0.0309 | 35.229 |
| Parsi_India | Parsi_Pakistan | Iranians | 0.0318 | 42.175 |
| Parsi_India | Gujaratis | Sindhi | -0.0015 | -1.89 |
| Parsi_Pakistan | Parsi_India | Sindhi | 0.0309 | 34.184 |
| Parsi_Pakistan | Parsi_India | Iranians | 0.0313 | 39.061 |
| Parsi_Pakistan | Gujaratis | Sindhi | -0.002 | -2.482 |
| Sindhi | Parsi_India | Parsi_Pakistan | 0 | -0.036 |
| Sindhi | Parsi_India | Gujaratis | -0.0038 | -4.609 |
| Sindhi | Parsi_Pakistan | Gujaratis | -0.0038 | -4.688 |
| Sindhi | Iranians | Parsi_Pakistan | -0.0124 | -16.085 |
| Sindhi | Iranians | Parsi_India | -0.0124 | -15.308 |
| Sindhi | Iranians | Gujaratis | -0.016 | -17.608 |
| Gujaratis | Parsi_Pakistan | Parsi_India | -0.0005 | -0.898 |
| Gujaratis | Iranians | Parsi_Pakistan | -0.0154 | -21.245 |
| Gujaratis | Iranians | Sindhi | -0.0211 | -22.669 |
| Gujaratis | Parsi_India | Sindhi | -0.0054 | -5.982 |
| Gujaratis | Parsi_Pakistan | Sindhi | -0.0058 | -6.7 |
| Iran_Neolithic | Sindhi | Iranians | -0.0002 | -0.141 |
| Iran_Neolithic | Gujaratis | Iranians | -0.008 | -5.075 |
| Iran_Neolithic | Parsi_India | Iranians | 0.005 | 3.2 |
| Iran_Neolithic | Parsi_Pakistan | Iranians | 0.0058 | 4.083 |

D= (Gp1,Yoruba;Gp2,Gp3)

We computed a maximum likelihood (ML) tree and co-ancestry matrix based on the haplotype structure of the Parsi populations, applying the default settings of ChromoPainter and fineSTRUCTURE (version1) (39). The ML tree split South Asian and West Eurasian populations into two distinct clusters (Additional File 1: Fig. S8). All the Parsi individuals form a unique sub-cluster embedded within the major West Eurasian population trunk. The co-ancestry matrix plot clearly differentiated Parsis from their neighbours in sharing a large number of chunk counts with West Eurasian (mainly Iranian and Middle Eastern) populations (Additional File 1: Fig. S9 and Supplementary text).. Additionally, South Asian populations have donated a significantly higher number of chunks to Parsis than they received from them (two tailed p value <0.0001). However, the number of these chunks was significantly lower than the chunk counts shared between any pair of South Asian populations (two-tailed p value <0.0001) (Additional File 1: Fig. S9).. The fineSTRUCTURE and *D* statistic results thus largely suggest unidirectional minor gene flow from South Asians to Parsis (Table 2 and Additional File 1: Fig. S9)..

To further investigate the parental relatedness among Parsis (19), we analysed the Runs of Homozygosity (RoH) and Inbreeding coefficient in the population (Additional File 1: Fig. S10). For RoH calculations, we applied three window sizes (1,000 kb, 2,500 kb and 5,000 kb), requiring a minimum of 100 SNPs per window and allowing one heterozygous and five missing calls per window. Long RoH segments characterise consanguinity and also provide a distinctive record of the demographic history for a particular population (40,41). As expected, both of the Parsi populations carried a larger number of long segments relative to their putative parental populations and present neighbours at the 1,000 kb window length, likely due to small population size and a high level of inbreeding. However, the Sindhi population from Pakistan also showed a higher level of inbreeding at the larger RoH window sizes, most likely due to an elevated level of cross-cousin marriages (Additional File 1: Fig. S10).

We used ALDER, a method based on linkage disequilibrium (LD), (42), to estimate the time of admixture between Parsis and their neighbouring South Asian populations. For this analysis we used the present-day Iranian *vs* Gujarati or Sindhi populations as their surrogate ancestors. We estimated the admixture time of Parsi groups to be around ∼ 40 (95% CI 26-50) generations, which yields a time of 1160 years (assuming a generation time of 29 years), in good agreement with their historically-recorded migration to South Asia (Table 3). We also tested evidence for a more complex admixture history using MALDER (43) which can be used to infer multiple admixture events. The MALDER analyses confirmed the ALDER results, demonstrating only one admixture event 54 ± 8 generations ago. The ancestral sources with the highest amplitude in MALDER were Sardinian and Dai (Additional File 1: Table S7).

**Table 3.** The formal text of admixture using ALDER method

| Reference 1 | Reference 2 | Admixed | Gen. Time | p value | Z score |
| --- | --- | --- | --- | --- | --- |
| Iranian | Gujarati | Parsi_India | 38.26 ± 12.16 | **0.0017** | **3.15** |
| Iranian | Sindhi | Parsi_India | 32.96 ± 9.42 | 0.013 | 2.48 |
| Iranian | Sindhi | Parsi_Pakistan | 41.32 ± 8.93 | **1.7x10<sup>-5</sup>** | **4.3** |
| Iranian | Gujarati | Parsi_Pakistan | 30.74 ± 14.04 | 0.029 | 2.19 |

To investigate how drift shaped the functional genetic variation after admixture in the Parsis, we implemented the population branch statistic (44) using the Sindhi and Iranians as reference and outgroup. We analysed variants over the 99.9th percentile of the genomic distribution, focussing only on those that were annotated as missense, stop gain, stop loss, splice acceptor and splice donor using the Ensembl Variant Effect Predictor tool (45). This revealed a cluster of linked SNPs in the HLA region and a missense SNP in CD86 (rs1129055) with a high ancestral G allele frequency in the Parsi (0.87) (Additional File 1: Fig. S11 and Table S8). The frequency of this G allele is lower in the Iranians and Sindhi (0.60) and other South Asians (0.52) and East Asians (0.40). This polymorphism has been associated with the pathogenesis of pneumonia-induced sepsis and the G allele has been associated with a decreased risk of active brucellosis in Iranians (46). The G variant has also been associated as an eQTL for decreased expression of *IQBC1*, an IQ motif containing B1 gene that is highly expressed in EBV transformed B lymphocytes (45).

To obtain a detailed understanding on the sex-specific South Asian and Iranian ancestries, we examined maternally-inherited mtDNA and paternally-inherited Y chromosome biallelic markers in a larger sample in both the Indian and Pakistani Parsi populations (Additional File 1: Fig. S12 and Tables S9 and S10). For the mtDNA analysis we were also able to assay 21 ancient samples from the Sanjan (21) region, for 108 diagnostic polymorphisms (Additional File 1: Table S11 and Supplementary text). Interestingly, we observed 48% South-Asian-specific lineages (haplogroups M2, M3, M5 and R5) among the ancient Parsi samples, which could potentially be explained in two ways. First, these haplogroups might have been carried by the migration of Zoroastrian refugees from Faras (Iran), a possibility which is supported by the presence of these clades in present-day Persian samples (9.9%) (34). Secondly, they might have resulted from the assimilation of local females during the initial settlement. The comparison of ancient and modern samples thus identified maternal lineages which can be considered as founding (surviving or lost), as well as those which were assimilated subsequently (Additional File 1: Fig. S12 and Tables S9-11). The Y chromosome profiles of Indian and Pakistani Parsi populations revealed a higher frequency of Middle-Eastern-specific lineages than South Asian ones in the Parsis (Additional File 1: Fig. S12 and Table S10). The PC analysis of both mtDNA and Y chromosome data placed all the Parsi groups close to each other and showed their contrasting clustering based on maternal or paternal ancestries (Fig. 4). For mtDNA, the Parsi cluster was closer to the Indian and Pakistani cluster (Fig. 4a), whereas for the Y chromosome it aligns between the Iranian and Pakistani populations (Fig. 4b). The Y-chromosomal PCA is similar to the autosomal PCA (Fig. 2 and Additional File: Fig. S1). The contrasting patterns of maternal and paternal ancestry support a largely female-biased admixture from the South Asian populations to the Parsis.

**Fig. 4.**
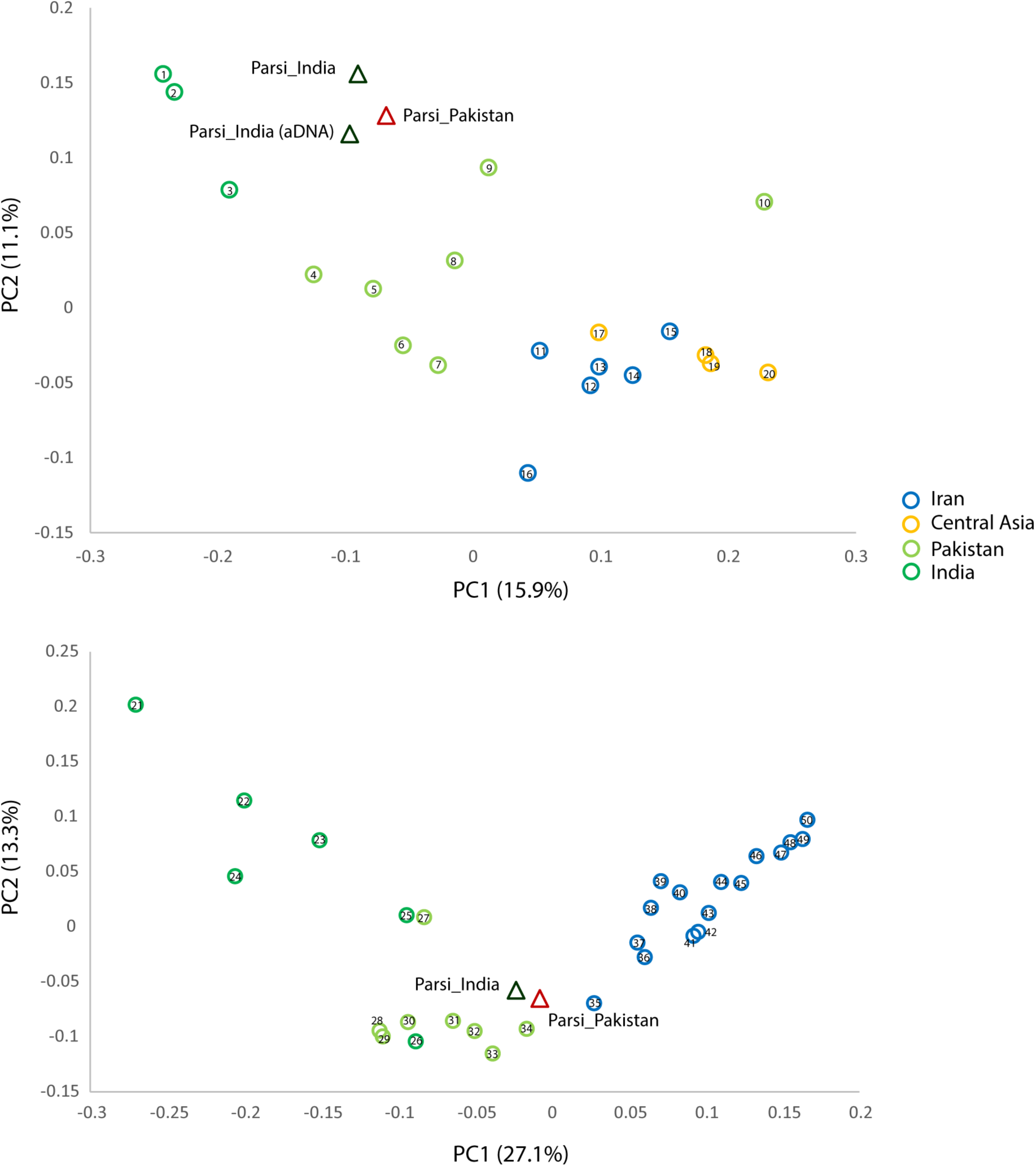
Principal component analysis using haplogroup frequencies for a) mtDNA, and b) Y-chromosome, **b)** in Parsis, Iranian, Central Asian, Pakistani and Indian populations. The details of populations is given in supplementary text.

## Conclusions

In conclusion, our investigation has not only contributed substantial new data, but has also provided a more comprehensive insight into the population structure of Parsis and their genetic links to Iranians and South Asians. We show that the Parsis are genetically closer to Middle Eastern (Iranian and Caucasian) populations than those in South Asia and provide evidence of sex-specific admixture with prevailing female gene flow from South Asians to the Parsis. Our results are consistent with the historically-recorded migration of the Parsi populations to South Asia in the 7th century and in agreement with their assimilation into the Indian sub-continent’s population and cultural milieu “like sugar in milk”.

## Methods

A detailed description of material and methods can be found in the supplementary text (Additional File 1). The Ethics Committee of the participating institutions (EBC, Estonia, CCMB, India and Human Materials and Data Management Committee, Wellcome Trust Sanger Institute) approved the study. The modern Parsi samples were pooled from three independent collections: two from Mumbai, India, and one from Karachi, Pakistan (Fig. 1). Illumina 650K and 2.5M chips were used to genotype 19 Indian and 24 Pakistani Parsi individuals, respectively, following the manufacturers’ specifications. We merged our newly-generated data of 43 samples with the relevant reference datasets of 829 samples published elsewhere (Additional File 1: Supplementary text and Table S2). For mtDNA control and coding region markers, we genotyped 117 Indian and 50 Pakistani Parsi samples (Additional File 1: Table S9). We followed phylotree (build 17) to classify them into haplogroups. For Y chromosome genotyping, 90 Pakistani samples were genotyped either by sequencing or by PCR-RFLP for the relevant Y chromosome markers, whereas 84 Indian samples were assayed for 80 Y chromosomal SNPs by using Sequenom mass array technology (Additional File 1: Table S10).

Ancient DNA samples were excavated from Sanjan, Gujarat in 2001 (Additional File 1: Supplementary Text). Archaeological analysis and AMS dating were consistent with these remains belonging most likely to the migrant Parsis during 8-13th Century (Supplementary text). The teeth obtained from 21 of these specimens were brought to the ancient DNA (aDNA) laboratory of the CSIR-Centre for Cellular and Molecular Biology, Hyderabad, India for the extraction and genotyping. We followed our standard published protocol to isolate DNA from the teeth (47) (Additional File 1: Supplementary text).

For autosomal analyses, after data curation and merging (Additional File 1: Supplementary text), we first used method of Cockerham and Weir (48) to estimate mean pairwise *F*st. Further, we performed PC analysis on pruned data using the smartpca v. 7521 (28) program (with default settings). We also used the *F*_*ST*_:Yes method of smartpca to calculate the *F*_*ST*_ with standard errors. We ran unsupervised ADMIXTURE v.1.23, 25 times each for K=2 to K=12, and used the method described previously to choose the best K value (31). The F statistics were performed by the ADMIXTOOL package v.3 (35) and the haplotype-based analysis was performed by Chromopainter and fineSTRUCTURE v.1 (39). The ML tree of world populations was constructed using TreeMix v.1.12 (38) and the RoH were calculated using PLINK 1.9 (49). ALDER v 1.03 (42) and MALDER v.1.0 (43) were used to calculate the time and number of admixture events. The population branch statistic method (PBS) (44) was used to identify genomic regions under selection in the Parsi population.

## Declarations

### Acknowledgements and funding

We are thankful to Dr. Shernaz Cama, Director Parzor Foundation, India for her help and critical comments. Tuuli Reisberg for technical assistance. Support was provided by: Estonian Personal grants PUT-766 (GC, MK, AKP); EU European Regional Development Fund through the Centre of Excellence in Genomics to Estonian Biocentre and Project No. 2014-2020.4.01.15-0012, and Estonian Institutional Research grants IUT24-1 (RV, MM, SR, EM, MR, TK); The Council of Scientific and Industrial Research (CSIR), Government of India (KT); Wellcome Trust grant 098051 (QA and CTS); WZCF (World Zarathushti Cultural Foundation), Parsi foundation and IAS, Indian Archaeological Society (VMT); PIRSES-GA-2012-318979 grant (MK). ERC Starting Investigator grant (FP7 -261213) (T.K.). AKP was supported by the European Social Fund’s Doctoral Studies and Internationalisation Programme DoRa. The funders had no role in study design, data collection and analysis, decision to publish, or preparation of the manuscript. The analyses were performed in the High Performance Computer Center of the University of Tartu, Estonia and the Wellcome Trust Sanger Institute, Hinxton, UK.

### Data availability

The data are available at NCBI GEO Accession no GSE97086 and the data repository of the Estonian Biocentre: www.ebc.ee/free_data.

### Author contributions

GC, QA, RV, KT and NR conceived the project and designed the experiments. GC, QA, NR, SP, AKP, RT, SF, MR, MK, DSR, AGR, EM, SR, and SK carried out haploid DNA genotyping and analysis. GC, QA, MaM and TK analysed autosomal data. VMT, KD and SRW performed archaeological study. VMT and NR collected ancient samples. NR isolated DNA from ancient remains and performed genotyping. SQM, LS, MM, CTS, RV and KT provided samples and reagents. GC, QA, NR, VMT and MaM wrote the manuscript with the help of TK, CTS, RV, MM and KT.

### Competing interests

None.

## Legends Additional files

**Additional File 1:** Supplementary text explaining the archaeological details of ancient samaples; isolation of ancient DNA, genotyping and statistical analyses. 12 figures and 11 tables are also incorporated in this file.

